# Methods in field chronobiology

**DOI:** 10.1101/197541

**Authors:** Davide Dominoni, Susanne Åkesson, Raymond Klaassen, Kamiel Spoelstra, Martin Bulla

## Abstract

Chronobiological research has seen a continuous development of novel approaches and techniques to measure rhythmicity at different levels of biological organization from locomotor activity (e.g. migratory restlessness) to physiology (e.g. temperature and hormone rhythms, and relatively recently also in genes, proteins and metabolites). However, the methodological advancements in this field have been mostly and sometimes exclusively used only in indoor laboratory settings. In parallel, there has been an unprecedented and rapid improvement in our ability to track animals and their behaviour in the wild. However, while the spatial analysis of tracking data is widespread, its temporal aspect is largely unexplored. Here, we review the tools that are available or have potential to record rhythms in the wild animals with emphasis on currently overlooked approaches and monitoring systems. We then demonstrate, in three question-driven case studies, how the integration of traditional and newer approaches can help answer novel chronobiological questions in free-living animals. Finally, we highlight unresolved issues in field chronobiology that may benefit from technological development in the future. As most of the studies in the field are descriptive, the future challenge lies in applying the diverse technologies to experimental set-ups in the wild.

## Introduction

For all organisms, exact timing of behaviour to both daily and seasonal environmental cycles is crucial for survival and successful reproduction [1,2]. Consequently, the study of biological rhythms, chronobiology, is a vibrant and interdisciplinary research area in biology [3–5]. However, chronobiology has been largely dominated by studies of just a few model organisms under standardized laboratory conditions [4]. Bringing such studies into the wild has often generated surprising outcomes [6–8].

The knowledge gaps and discrepancies between laboratory and field studies were emphasized in a recent perspective article on the diversity of animal clocks in the wild: “…*to begin to understand the adaptive significance of the clock, we must expand our scope to study diverse animal species from different taxonomic groups, showing diverse activity patterns, in their natural environments*” [4]. Indeed, whereas controlled laboratory studies are essential to investigate the proximate mechanisms behind biological rhythms, they offer little insight about the diversity of temporal strategies that free-living animals may adopt and the fitness consequences of an eco-evolutionary process that takes place in the “real world”.

Building on this evidence, ecologists are increasingly using individual-based telemetry with high temporal and spatial resolution not only to study the movements of wild animals, but also to gain insights into the temporal patterns of their behaviour and physiology, as well as into the genetic, environmental and/or life-history factors that might affect the regulation of such rhythms [9,10]. For instance, recent telemetry work on arctic shorebirds has revealed unexpected inter- and intra-specific variation in their behavioural rhythms under the continuous daylight of arctic summer [10–12], and the combination of automatic radiotelemetry and EEG loggers demonstrated a link between activity and fitness in a polygynous shorebird [13]. Also, combining GPS-tracking, accelerometers and EEG loggers revealed unprecedented sleep-wake cycles of frigatebirds (*Fregata minor*) flying over the ocean for up to 10 days uninterruptedly [14]. The integration of ecological and chronobiological approaches and techniques can therefore help not only answering old questions traditionally confined to laboratory settings, but also ask novel exciting questions that are more pertinent to field systems (Table 1). Such integration can also improve our understanding about how adaptive biological rhythms are in the wild, for instance through a combination of genetic engineering and animal tracking [15].

**Table 1.** Examples of pressing questions in the field chronobiology.

| # | Question | Case study |
| --- | --- | --- |
| 1 | How variable are rhythms within and between taxa, species, populations and individuals? | 1, 2 |
| 2 | What drives such variation? (e.g. environmental conditions, internal state, sociality, anthropogenic disturbance, global environmental change) | 1, 2 |
| 3 | Are rhythms of captive animals comparable to rhythms of animals in the wild? | 1 |
| 4 | How can we disentangle the relative contribution of endogenous (genes) versus exogenous (environment) drivers of rhythmicity in wild species? |  |
| 5 | Is variation in rhythms associated to fitness? | 3 |

This paper has two major aims. First, we review traditional and relatively recent tools to collect chronobiological data in the wild. In particular, we emphasize the suitability of ecological tools initially developed for other purposes (e.g. to map migration patterns of animals) and the added value of integrating different technologies. Second, we present three case studies to demonstrate how such tools and their integration can be used to answer some of the questions in the field chronobiology (Table 1). We pick each case study for a specific reason. The first case study uses array of technologies to reveal the diversity and drivers of behavioural rhythms in the wild, as well as to discuss how the findings compare to the findings from captive conditions (Table 1, questions 1-3). Most of the technologies in this case study were traditionally deployed for other purposes than measuring behavioural rhythms. In the second case study we initially review technologies that allow individual-based and group-based tracking of insects, for which we have yet to fully appreciate how their rhythms are expressed in the wild. We then demonstrate the use of laser radar to record activity rhythms of insect groups across time and habitats (Table 1, question 1). The third case study combines tracking methods and genetic engineering to tackle one of the most pressing chronobiological questions, that is, whether clocks are adaptive (Table 1, question 5).

## Integrating old and new approaches to record rhythms in the field

Chronobiologists assess rhythmicity in captive animals by measuring activity rhythms (e.g., locomotion and foraging), physiological rhythms (e.g., body temperature or melatonin production), and molecular rhythms (e.g., gene expression) [3,16,17]. Activity rhythms are quantified using infrared sensors or mechanical instruments such as the running wheel [18]. Physiological rhythms are usually assessed using temperature and heart rate loggers [19], or sampling of blood, urine and faeces which are subsequently analysed for hormone concentration (melatonin, testosterone, etc.) [20]. Molecular rhythms are assessed by gene expression - a relatively recent tool - performed with diverse methods ranging from microarrays to quantitative PCRs [17] and transcriptomics [21], or by quantification a wide range of proteins and metabolites [21]. All these methodologies can be used, and some of them already are used, to also elucidate rhythms of organisms in the wild. For instance, a recent study used running wheels with free-living mice in the wild [18] and found similar temporal patterns of running as in captive mice. In addition, there have been great developments in individual-based tracking technologies as well as in automated monitoring systems, which allows gaining unprecedented insight into behavioural and physiological rhythms of free-living animals. Thus, chronobiologists have now a well-equipped toolbox at hand to study rhythms of organisms in the wild.

We summarise the methods available to field chronobiology in Table 2 and 3. We distinguish methods used to record behavioural and physiological rhythms (Table 2), which often involve tagging animals, from relatively new methodologies that assess molecular rhythms or use genetic engineering to manipulate circadian time (Table 3). We briefly describe how each method works, what kind of rhythmic information it can measure, and provide examples of chronobiological questions it can help answer. Although we have described each method separately, field chronobiology may strongly benefit from integrating existing methodologies. For instance, geolocators and accelerometers can be jointly deployed on the same animal to infer daily activity patterns of birds at different stages of their migration journey [22,23]. and a combination of accelerometers, automated radio-telemetry and EEG recordings revealed strikingly variability in timing of sleep in tree-toed sloths (*Bradypus variegatus*) [24]. In addition, different technologies can be integrated within a single tag. For example, daily diaries have multiple built-in sensors that simultaneously record behavioural, physiological and environmental rhythms [25], thereby allowing a holistic view into biological rhythms of wild animals.

**Table 2.**
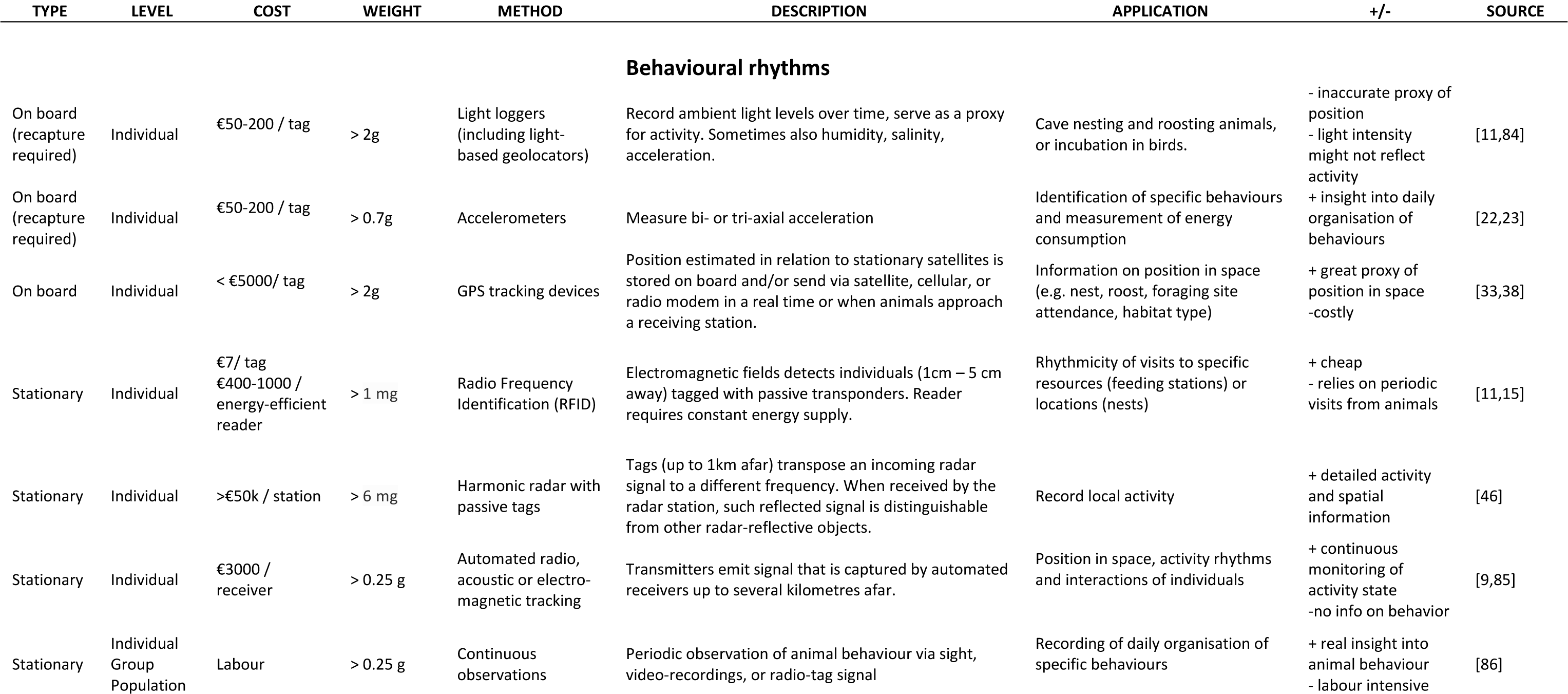
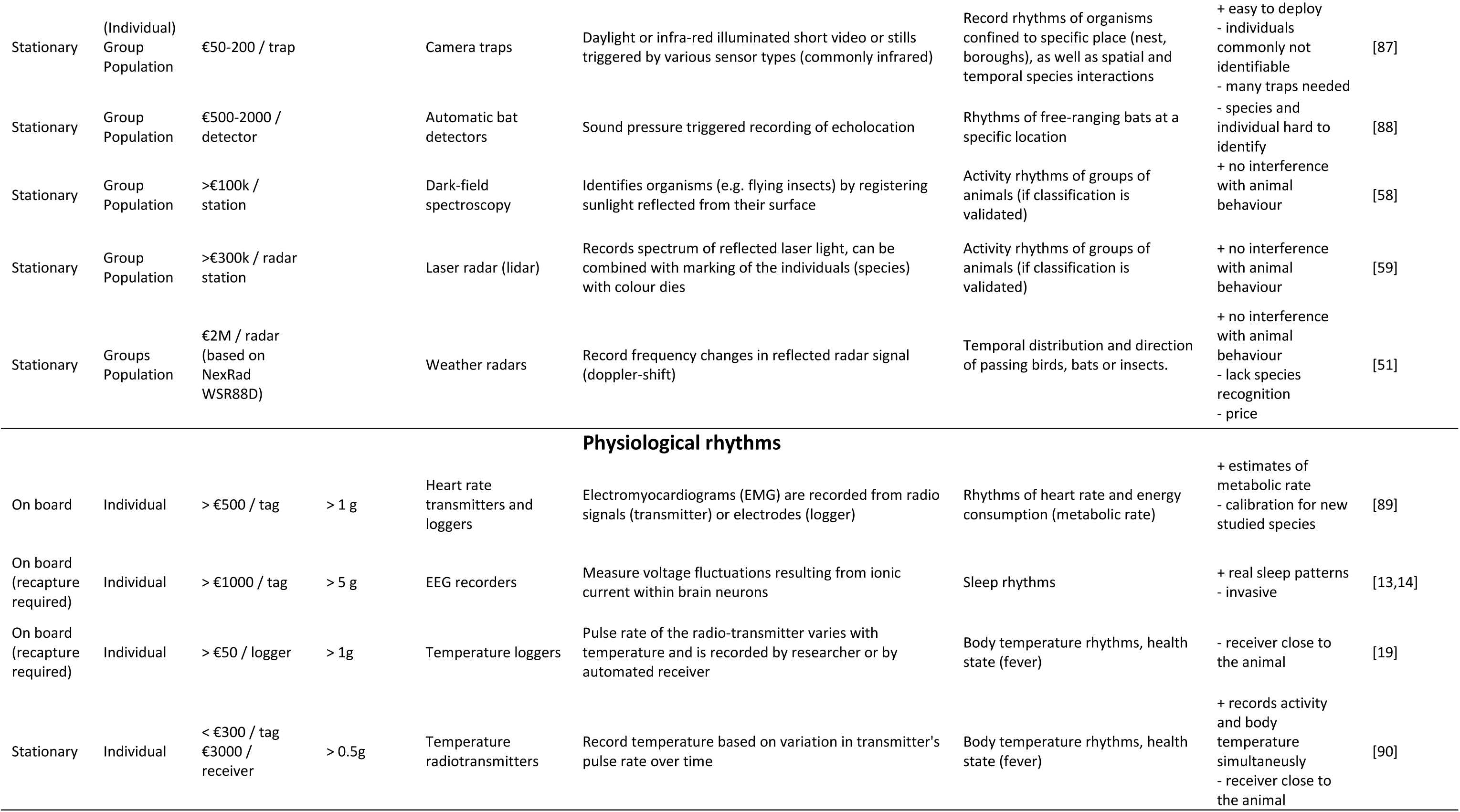
Methods to record behavioural and physiological rhythms in free-living animals. Methods ordered according to (1) type, (2) level of application, (3) tag weight and (4) price. Specific applications are listed only when unique to the given method. For instance, for behavioural rhythms all methods are applicable to record activity vs. non-activity and hence that is not stated.

## CASE STUDY 1: Diversity of individual rhythms in the wild - avian incubation and foraging behaviour

*Key questions*: How variable are rhythms within and between wild populations? What drives such variation? Are rhythms of captive animals comparable to rhythms of animals in the wild?

We understand little about within- and between-species diversity of behavioural rhythms in the wild (see Editorial of this issue and [4]). Consequently, we also understand little about what drives the potential variation in these rhythms, e.g. to what extent rhythms are determined by evolutionary history and/or by plastic responses to the environment.

Here, we demonstrate how diverse monitoring methods can be used to fill this knowledge gap, that is, to study variation in behavioural rhythms (daily and seasonal) in free-living non-model organisms in their natural environments and in unexplored contexts. Specifically, we discuss monitoring methods used to reveal the diversity in incubation rhythms of biparental shorebirds [11] and demonstrate the use of GPS-tracking to derive novel data on diverse foraging activity patterns of raptors (Klaassen *et al*., in preparation).

### Incubation rhythms of biparental shorebirds

It is often unclear how findings on single individuals translate to the social context typically experienced by organisms in their natural environment, i.e. when it matters to them [4]. For example, when individuals pursue a common goal such as reproducing, the social environment is expected to shape their behavioural rhythms [26]. However, social synchronization and its outcome in terms of behavioural rhythms are poorly understood.

Avian biparental incubation is a mutually exclusive, but socially synchronized, behavioural rhythm. A recent study [11] used an array of monitoring methods (RFIDs, light loggers, GPS-based systems, radio-tags, video recordings and continuous observations; Table 4) to reveal unprecedented within- and between-species diversity in incubation rhythms across 729 nests of 91 populations of 32 biparentally-incubating shorebird species (Fig. 1).

**Figure 1.**
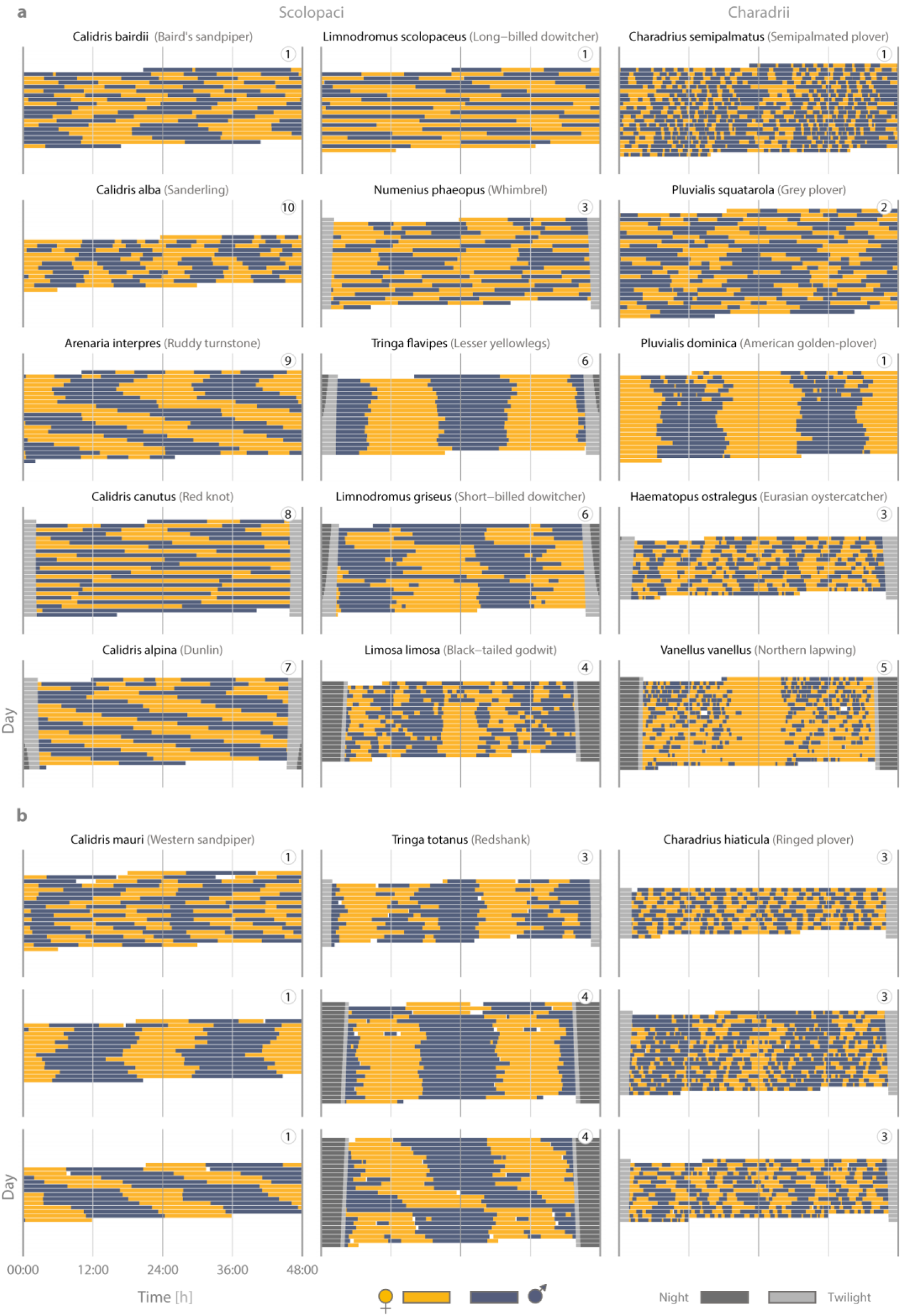
Actograms illustrating the diversity of shorebird incubation rhythms. **a**-**b**, Each actogram depicts the bouts of female (yellow; 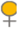) and male (blue gray;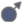) incubation at a single nest over a 24-h period, plotted twice, such that each row represents two consecutive days. If present, twilight is indicated by light grey bars 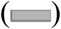 and corresponds to the time when the sun is between 6° and 0° below the horizon, night is indicated by dark grey bars 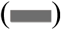 and corresponds to the time when the sun is > 6° below the horizon. Twilight and night are omitted in the centre of the actogram (24:00) to make the incubation rhythm visible. The circled numbers 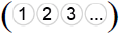 indicate the breeding site of each pair (i.e. highlight which pairs bred in the same breeding site). **a**, Between-species diversity. **b**, Within-species diversity. Note that the three rhythms for Western sandpiper and Ringed plover come from the same breeding location. The actograms for each nest in the study together with the data and code to replicate the figure are freely available at <u>https://osf.io/wxufm/</u> [28]. This figure was adopted from [11].

Multiple sampling methods allowed us to include more species and populations, as well as to increase sample size for some populations. Although the sampling interval varied from continuous to 30 min sampling between methods (Table 3) and populations, as well as within some populations, the incubation variables were independent of sampling interval (Table 2 in the Extended Data of [11]).

**Table 3.** Methods to record molecular rhythms in free-living animals.

| METHOD | DESCRIPTION | APPLICATION | COST | + | - | SOURCE |
| --- | --- | --- | --- | --- | --- | --- |
| Fibroblast cells | A clock gene probe flagged with a bioluminescent marker is inserted into fibroblasts cultured from skin biopsy. Rhythmic expression of clock gene is then measured in constant darkness in a lumicycle. | Measure circadian period length in fibroblast (dermal) cells | > €100 per sample (after first investment to set up facilities) | - Insight into endogenous circadian traits<br>- Useful to disentangle endogenous (clock-driven) from environmental factors affecting rhythmicity | - Cell cultures facilities needed<br>- Requires the use of a lumicycle (> €50k) | [41] |
| Gene expression | Gene expression level is quantified via transcriptomics, microarrays, RT-qPCR | Diel and seasonal expression levels of candidate genes as well as of entire gene pathways (e.g. immunity, antioxidant system etc) | > €200 / sample | - Expression levels of candidate genes involved in timing, also thousands of genes simultaneously (transcriptomics) | - Bioinformatics<br>- Reference genome | [21] |
| Proteomics/<br>Metabolomics | Relative abundance of proteins and metabolites is quantified via mass-spectrometry | Diel and seasonal abundance of proteins and metabolites (including hormones) | > €200 / sample | - Allow studying the abundance of hundreds of proteins and metabolites simultaneously | - Bioinformatics | [91] |
| Genetic engineering | e.g. mutagenesis, gene targeting by homologous recombination, gene replacement | Daily and seasonal rhythms in sleep, activity, physiology in mutant animals lacking key circadian genes | €5000-10000 (for mice, less predictable for other animals) | - Test properties and consequences of specific aspects of timing | - Pleiotropic effects may affect other genes than target circadian ones | [15] |

**Table 4.** Methods used to derive incubation in biparental shorebirds

| Method* | How was incubation derived? | *Sampling interval*<br>*(min)* | *N*<br>*populations* | *N*<br>*nests* |
| --- | --- | --- | --- | --- |
| RFID with temperature probe between the eggs | A thin antenna loop, placed around the nest cup and connected to the reader, registered presence of tagged parents at the nest; the passive-integrated tag was either embedded in a plastic flag [12] with which the parents were banded, or glued to the tail feathers [42]. Temperature recordings allowed to identify whether a bird was incubating even in the absence of RFID readings; an abrupt change in temperature demarcated the start or end of incubation [12]. | 0.08-5.5 | 23 | 200 |
| light logger attached to bird's leg | The logger recorded maximum light intensity for a fixed sampling interval (2-10 min) and recorded darkness when parent was incubating. An abrupt change in light intensity (as opposed to a gradual change caused, e.g. by civil twilight) followed by a period of low or high light intensity demarcated the start or end of the incubation period [11]. | 2-10 | 71 | 396 |
| GPS tag mounted on the back of the bird | The tag recorded the position of the bird [43] and incubation was assumed whenever the bird was within 25 m of the nest. | 10-30 | 2 | 9 |
| Automated receivers | Receivers recorded signal strength of a radio-tag attached to the rump of a bird; whenever a bird incubated, the strength of the signal remained constant [10]. | 0.07 | 2 | 3 |
| Video cameras and continuous observations | Videos and observations were used to identify the incubating parents; parent identification was based on plumage, colour rings or radio-tag. | constant - 30 | 6 | 61 |

\* For technical specifications of the methods see Extended Data Table 1 in [11].

Incubation records were transformed to local time (UTC time+(nest's longitude/15)) to make them comparable across sites. For each nest, the authors manually or automatically [11,12,27,28] extracted lengths of all available incubation bouts defined as the total time allocated to a single parent (i.e. the time between the arrival of a parent at and its departure from the nest followed by incubation of its partner). Bout lengths were then used to extract the length of the period (the most prominent cycle of female and male incubation) that dominated each incubation rhythm. Finally, phylogenetically informed comparative analyses were used to investigate phylogenetic signal in bout and period length, the relationship between bout length and body size, latitude and escape distance from the nest, as well as relationship between period and latitude.

The study found substantial within- and between-species variation in incubation rhythms (Fig. 1). For example, between species, the period length of the incubation rhythms varied from six to 43 hours. Different species, but also different pairs of the same species, adopted strikingly different incubation rhythms, even when breeding in the same area. For example, the incubation period length for Long-billed dowitchers *Limnodromus scolopaceus* varied from 21.75 to 48 hours. Interestingly, 24-h incubation rhythms were absent in 78% of nests representing 18 out 32 species.

Importantly, the study explained part of the described variation in the incubation rhythms. For example, there was a strong phylogenetic signal (Pagel’s λ was close to 1). In addition, the incubation rhythms with periods that do not follow the 24-h light-dark cycle were more common and the deviations from 24-h increased in shorebirds breeding at high latitudes. This supports the existence of a latitudinal cline in incubation rhythms, but a substantial number of rhythms defied the 24-h day even at low and mid latitudes. These results indicate that under natural conditions social synchronization can generate far more diverse behavioural rhythms than previously expected (e.g. from studies of captive animals), and that the incubation rhythms often defy the assumptions of entrainment to the 24-h day-night cycle.

### Diel activity patterns of diurnal raptors

Individual variation in daily and seasonal foraging rhythms remains poorly understood. This is perhaps not surprising as, until recently, long term monitoring of many individuals was not feasible (e.g. it was too labour intensive, but see an example on hunting activity of individual European Kestrels *Falco tinnunculus* recorded with visual observations [29]). This issue is now solved by the availability of several types of tracking devices that allow us to follow the behaviour and movements of individual animals in unprecedented spatiotemporal detail (Table 2). However, most analyses of tracking data focus on spatial aspects such as home range size and migration routes, whereas temporal aspects such as daily and seasonal activity patterns are largely overlooked (but see e.g. [30–33]. This suggests that the huge amount of detailed tracking data that is currently routinely collected is generally underused for chronobiological purposes. Here we provide an example of how GPS-tracking data could be used to infer daily foraging rhythms of individual Montagu’s Harriers *Circus pygargus*.

We re-analysed GPS tracking data of three individual Montagu's Harriers, which were originally collected to study home range behaviour and habitat use during the breeding season (Klaassen *et al*. in preparation). The birds were tracked by UvA-BiTS GPS-loggers [34] that were programmed to sample the position and speed of the bird every 5 to 30 minutes during the day and every hour to two hours during the night (note that one of the advances of this tracking system is that tags can be programmed remotely when in reach of a local antennae, [34]). Flying behaviour was defined when the instantaneous GPS-speed exceeded 2 m/s, as the intercept of the probability density functions of speeds during sitting and during flight, which together make up the bi-modal density distribution of instantaneous speeds (Klaassen *et al*., in preparation). To reconstruct daily activity patterns, the proportion of flight instances within each hour of the day was calculated, lumping data across all available days (7-14 days, see Fig. 2). The average time flying per day was obtained by the sum of the hourly flight proportions. In this analysis, only daylight hours were included as loggers only collected sufficient data during daylight hours (fix interval 5-30 minutes) and because harriers are strictly diurnal. Montagu's Harriers hunt on the wing by slowly flying above foraging habitat, thus flight will mainly represent foraging activity. As Montagu's Harriers are long-distance migrants, daily activity patterns could be compared across different ecological contexts, i.e. the breeding site in Europe, their main migratory stopover site in Northwest Africa, and the wintering site in the Sahel in Africa [35].

**Figure 2.**
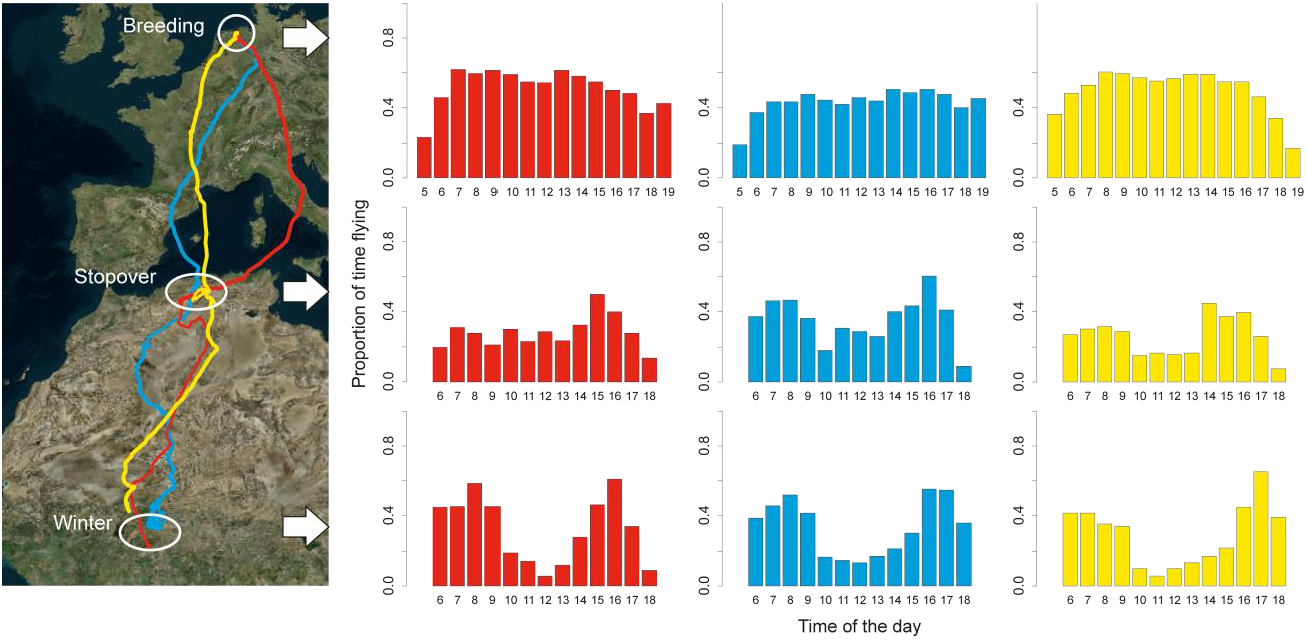
Example of daily foraging rhythms in three Montagu’s Harriers. Map presents migration tracks and corresponding bar plots represent proportion of time a bird spent flying during each hour of the day (GMT; averaged across 7-14 days) at the wintering site (lower plots), main spring migratory stopover site (middle), and breeding site (upper). Note that only daylight hours are included as Montagu’s Harriers are strictly diurnal and loggers recorded the necessary detailed information (5-30 min sampling interval) only during the day. The three harriers are distinguished by colour and name.

Within individuals, daily activity patterns differed between the breeding, stopover and wintering site. Harriers flew more at breeding sites (mean values for the three individuals were 6.4, 7.5 and 7.7 hours per day, based on N = 183 days) than at stopover (3.4, 3.7 & 4.7 hours per day, N = 42 days) or wintering sites (4.1, 4.2 & 4.4 hours per day, N = 214 days) (Fig. 2). Not only had the amount of time spent flying per day varied between sites, but also temporal patterns of daily activity. For example, at wintering sites, harriers have a distinct dip in activity around noon. This “Siesta” is much less pronounced for stopover sites and almost absent for breeding sites (Fig. 2). Activity patterns also differed between individuals, with for example “Joey” flying less (6.5 hours per day) during the breeding season compared to “Elzo” (7.7) and “Yde” (7.5) (Fig. 2).

In order to quantify the degree of similarity in daily activity patterns, for example between individuals or between sites, the overlap index was calculated:

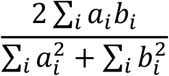

where a and b are the proportions of time flying for the two activity patterns that are compared, for different hours (i). This index ranges from 0 for non-overlapping distributions to 1 for identical distributions [36]. The overlap index between sites (average [range] index for each individual: red = 0.81 [0.77-0.87], blue = 0.90 [0.85-0.95] and yellow = 0.75 [0.73-0.79]) was relatively low compared to the overlap index between individuals at a given site (average index [range]: breeding = 0.97 [0.96-0.99], stopover = 0.94 [0.93-0.95] and wintering = 0.93 [0.88-0.96]). Thus, in this particular example, activity patterns tended to be similar across individuals at a given site, but varied to a greater degree across sites (Fig. 2), possibly suggesting a prominent role of environmental drivers shaping activity patterns in wild Montagu’s Harriers. Seasonal differences in daily activity patterns could, for instance, arise from different feeding habits of Montagu's Harriers in the three different seasons: voles and the need to feed young during the breeding season [37], eggs and nestlings of passerines during the main spring stopover in NW Africa [38], and grasshoppers in winter [39]. Whether between-individual variation in activity patterns reflects differences in individual personalities, with some birds being more explorative than others (e.g. [40]) or differences in habitat quality, is unclear. The speculations about how this within- and between individual variation arise deserve future testing and can only be resolved by combining the tracking results with field observations of the ecological circumstances at the three sites, such as how prey abundance and harriers' hunting success vary throughout the day [38].

### Conclusion

We demonstrated how incubation and foraging rhythms of free-living birds vary within- and between individuals and species, across seasons, latitudes, and depending on phylogeny (i.e. provide answers to question 1-2 in Table 1), and that such rhythms are more diverse than expected from studies in captivity (question 3 in Table 1).

These finding generate three main questions: (1) Are other behavioural rhythms in the wild also that diverse? (2) Are these rhythms regulated by endogenous (clock-driven) or environmental factors, or by a mixture of these? (3) What are the fitness consequences of various behavioural rhythms? To address these questions, we need to (1) expand our studies to different species and ecological contexts (e.g. monitoring of rhythms in both predators and preys), (2) use molecular tools that allow quantification of endogenous clocks in the wild (e.g. fibroblast assays, see Table 3 and [41,42]), and (3) monitor behavioural rhythms over the long-term to understand if and how individual variation in rhythms is linked to fitness (see case study 3).

## CASE STUDY 2: From individual to population rhythms: Timing of insects’ movement in the field

*Key questions*: How variable is timing of activity between insect groups? What environmental factors are related to such variation?

Insects are key laboratory models in chronobiology [43,44]. Yet, long term biotracking of insects in the wild, unlike tracking of vertebrates [45], is rare and limited to the largest species [46]. This is alarming because the limited evidence from semi-natural conditions revealed temporal components of behaviour that markedly differ from those recorded in the laboratory [8]. Here we briefly review the tracking of individual insects, as well as of groups (for description of each method). Then we illustrate recent applications of laser radar to identify groups of insects and their daily activity rhythms over various habitats.

### Tracking individual insects

Monitoring of individual insects in the wild can be done by active (battery-powered) radio transmitters or by harmonic radar and RFID which use passive tags (without battery) [46]. Radio-telemetry is limited by the available tags, most of which are too large, too heavy (2-100% of body mass), have limited tracking range on the ground (100–500 m), and/or have short battery life (7-21 days) [46]. Hence, radio-telemetry has been mainly used with larger insects (beetles and crickets), and only relatively recently with bees, dobsonflies and dragonflies. Such studies are mainly local in scale, but ground crews and receivers mounted on an airplane allowed monitoring of dragonfly migration over 150 km and up to 12 days [47].

In contrast to radio-transmitters, the tags used with harmonic radar and RFID have lower weight and hence can be used with a broader range of insect taxa [46]. Although the individuals can be monitored over a longer period of time than with radio-transmitters, the monitoring is only local as the detection zone of a stationary radar unit is < 1 km in diameter and the detection distance of RFID tags is usually < 1–5 m. Thus, RFID is useful for insects returning on a regular basis to their burrows (e.g. crickets) or hives (e.g. bees and bumblebees) [46,48].

Although miniaturization of tags will certainly extend the range of trackable insect taxa, some miniature insects are trackable only in groups, for instance with help of citizen science [49] or various radar technologies [50–52] (see Table 2 and next section).

### Tracking groups of insects

Vertical-looking radars, harmonic radars and weather radars have all been deployed to track flying insects since 1970s [50–52]. Vertical-looking radars detect insects that pass through the radar beam pointing up into the sky. Harmonic radars detect movements across a horizontal transect at a ground level, while the beam of weather radars spreads out as it moves away from the station, covering an increasingly larger volume (up to several km^3^). Thus, these radars are useful to infer timing of migration, flight altitudes (up to 1km) and orientation of the insects in relation to winds [53]. However, information about movements is generally limited to a single location of observation [54]. Moreover, this technology is suitable predominantly for large insects, and insects can only be classified by size and air speeds. In sum, these radar technologies are usually unable to distinguish species from one another. However, to understand activity rhythms in free-ranging insects, especially of those that are too small for any individual-based tracking technology, identifying insects remotely to groups, families, or better to species, is necessary.

Classification of insects to groups may be possible with laser radar (lidar; for details see below and [55]). The lidar beam that spreads out as it moves away from the station, covers a probe volume of approximately 10 m^3^ over a 2 km range. Lidars can detect groups of insect by measuring the spectrum of the light reflected by the body and wings of the flying insect as it flies across the laser beam [55,56]. That is, lidar can classify groups of insects according to wing beat frequency, body size, wing area, and potentially also body surface structures.

Classification of larger insects (such as damselflies) to species and to sex (if sexes are colour dimorphic) is also possible. Individuals previously marked with fluorescence dye generate a colour specific peak in the lidar signal [57]. Alternatively, dark-field spectroscopy identifies flying insects by registering sunlight reflected from the insect surface, when the insect passes across sampling area (ø 20-30 cm, up to 300 m afar) monitored by the spectroscope [58]. The distance to the insect is measured as well as its size and direction of flight, thereby including also spatial components of activity patterns.

### Use of laser radar to identify rhythms in groups of flying insects

Here, we illustrate the use of lidar technology to record temporal and spatial variation in flying insect abundance according to insect groups and habitat structures [59]. In this study, the lidar beam was sent 1.8 m above an open meadow and terminated at a distance of 140 m by a box made out of a dark cardboard, in an area where the meadow was surrounded by a forest edge.

Over the course of two nights, three main insect clusters were identified in the data (Fig 3a). Some insect groups had a wider peak of activity and were more evenly distributed over the 140 m transect (Fig 3b, Cluster a-b) than others (Fig 3b, Cluster c). Specifically, one insect cluster was especially abundant at the beginning of the night (Fig. 3b, Cluster c), especially in a meadow surrounded by a forest edge. Such temporal structuring across habitats might by typical for insects. For example, abundance of flying insects is higher over the grazed meadow compared to crop fields with oats [60].

**Figure 3.**
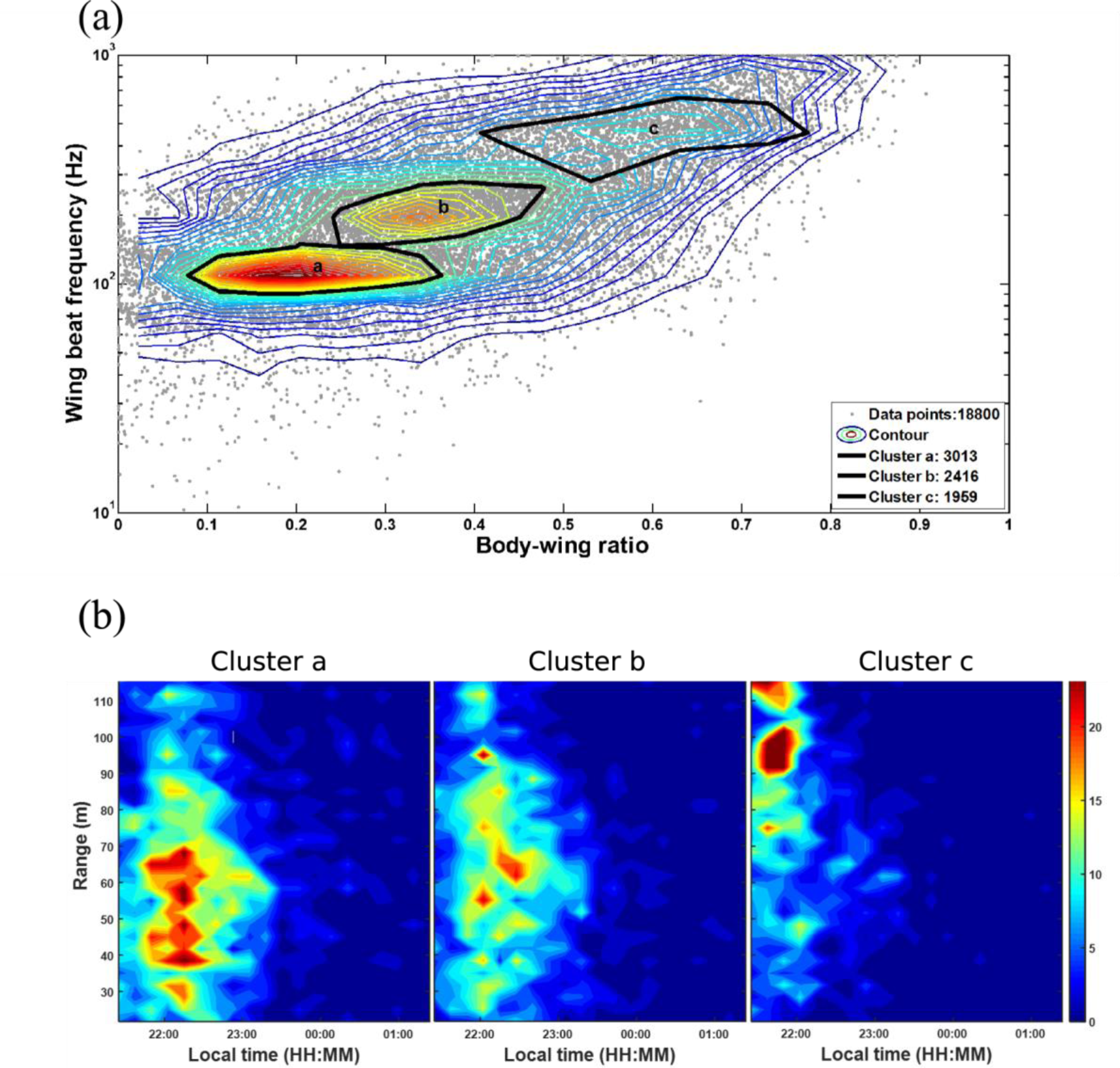
Using lidar to classify insect groups and their temporal and spatial distribution. (a) Contour plot illustrates insect densities based on body-wing ratio and the wing beat frequency as recorded by lidar. The three major insect clusters are indicated by black curves and as verified by insect traps represent mostly *Trichoptera* and *Chironomidae* (Cluster a), swarming non-biting midges and fies (Cluster b), and compact insects (Cluster c). The number assigned to each cluster in the legend represents the number of points in that cluster [59]. (b) Heat maps with temporal and spatial distribution of the three insect clusters. Note that the 140m long transect (range) started in an open meadow and terminated in a meadow surrounded by a forest edge. (a-b) Red denotes areas with a high density of insects, whereas blue denotes low insect densities. Image reproduced with permission from [59].

These findings elucidate how various insect groups cluster in time and space and suggest variability in daily timing across different groups of insects and across various habitats. However, the study has two major limitations. First, the study lasted only for three days, but recordings over several days, preferably months, are necessary to identify activity rhythms and their variation over time. This is essential, if we aim to elucidate the role of different environmental variables in driving variation in such rhythms. Second, the body-wing proportions overlapped between species and insects were thus classified only to groups based on body-wing proportions and wing-beat frequencies (Fig. 3a). However, deeper understanding of insect behavioural rhythms requires classification of insects down to the order, family or better species level. Such classification might be feasible if the species classification algorithm includes additional variables [60], or when it is calibrated by releasing insects of known species that are then recorded by the lidar.

### Conclusion

Here we briefly reviewed the technology and limitations to track insects, both individually and in groups. Then we demonstrated how lidar may reveal temporal and spatial variation in activity of various flying insect-groups [59,60]. Hence, the study provides preliminary insights about how insect rhythms vary between groups (Table 1, question 1) and across habitat types (Table 1, question 2). We further highlight current limitations in classifying insects to lower taxonomic levels. Once such limitations are tackled, lidar will help us answer question related to the variability and drivers of rhythms in different insect taxa, and how these differ between laboratory and wild populations (Table 1, question 3).

Lidar technology might also proof suitable for future investigations of nocturnal bird migration. Species could be classified based on flight speed, but also based on plumage characteristics, including coloration [57,61]. Such information is of interest to comparative studies investigating seasonal and diurnal variation in migration patterns. In addition, although lidar has been so far applied mainly in pilot studies over short time period, using this technology over longer periods will improve our understanding about daily rhythms of insect abundance across seasonal and environmental contexts.

## CASE STUDY 3: Measuring fitness consequences of circadian organization in the wild

*Key question*: Can we link the variation in circadian organization of activity to fitness in the wild?

Accurate timing of daily activity of organisms has long been assumed essential for fitness and survival, for example for the anticipatory regulation of physiology and behaviour in advance of changes in environmental conditions [62,63]. In captive animals, positive effects of a near 24 h endogenous circadian period with a duration comparable to the external (laboratory controlled) light-dark cycle are reported for the growth rate and longevity of insects [44,64], as well as for the lifespan of mice [65] and hamsters [66]. However, the laboratory is not the environment in which species have evolved and circadian traits have been selected. It is therefore essential to study the adaptive value of circadian function under natural conditions [67,68].

To demonstrate adaptiveness of circadian organization in natural habitats is however daunting. First, the powerful natural light/dark cycle limits experimental manipulation of temporal behaviour. Second, free-ranging individuals have to be followed throughout their life and their reproductive success needs to be measured. DeCoursey et al. [69,70] studied fitness consequences of lesions of the Suprachiasmatic Nucleus (SCN), the master circadian pacemaker in mammals, in antelope squirrels (*Ammospermophilus leucurus*) and chipmunks (*Tamias striatus*). SCN lesions are used in chronobiology experiments to abolish circadian rhythms in sleep-wake cycles and activity in mammals [71,72]. The survival of antelope squirrels and chipmunks was monitored with the use of transponders (RFID’s, see Table 2) and radio telemetry, respectively. In these studies, the SCN lesions compromised longevity: individuals with lesions lived shorter than sham control animals. These results provided the first evidence of the adaptive value of circadian organization in free-ranging mammals. However, to rigorously test for fitness consequences, it is essential to measure whether circadian rhythms not only affect survival, but also reproductive success [67].

A way to measure both individual and reproductive fitness is to use heritable circadian traits (e.g. the level of rhythmicity or the length of internal clock’s circadian period) in a selection experiment. Such traits have become available in an increasing number of organisms in the form of natural or engineered circadian mutants. Selection experiments have been done in the laboratory with strains of cyanobacteria carrying mutations that effected their circadian period. Strains with a circadian period similar to the applied external light-dark cycle outcompeted strains with a different circadian period; thus, showing selective advantage for an endogenous circadian period that matches the external light/dark cycle [73,74].

Here, we highlight the methods to translate selection experiments into semi-natural conditions using results from two competition experiments with mice [15,75]. The experiments integrated existing monitoring methods with present-day availability of circadian mutants. Wild-type mice (without the mutation) and mutant mice (homo- and hetero-zygote for a circadian mutation) were housed in mixed populations in outdoor enclosures. All mice were produced from heterozygote parents. Mice presence and longevity was monitored by subcutaneous RFID tags recorded at feeding stations. This allowed permanent monitoring of each individual and hence the mutant allele frequency in each population.

The first study [75] used mice with a mutation in the period2 gene (*mPer2*^*brdm*1^), which weakens circadian rhythmicity and causes health problems in the laboratory [76]. The mutant and wild type mice were released into four outdoor enclosures in near Mendelian ratio (homozygote: heterozygote: wild type = 1:2:1). However, there was no selection against the mutant allele over the course of two consecutive years [75].

The second study [15] used a comparable setup as the first one, but with six outdoor enclosures and the mutant tau allele (*Ck1e*^*tau*^) which shortens the endogenous circadian period [77]. At the start of the experiment this mutation was present in near Mendelian ratio. Here, a strong selective force against the mutant allele reduced its frequency from approximately 0.5 to almost 0.2 in little over a year (Fig. 4). Even though unknown non-circadian pleiotropic effects by the mutation cannot fully be excluded, this finding strongly indicates fitness consequences of aberrant circadian organization.

**Figure 4.**
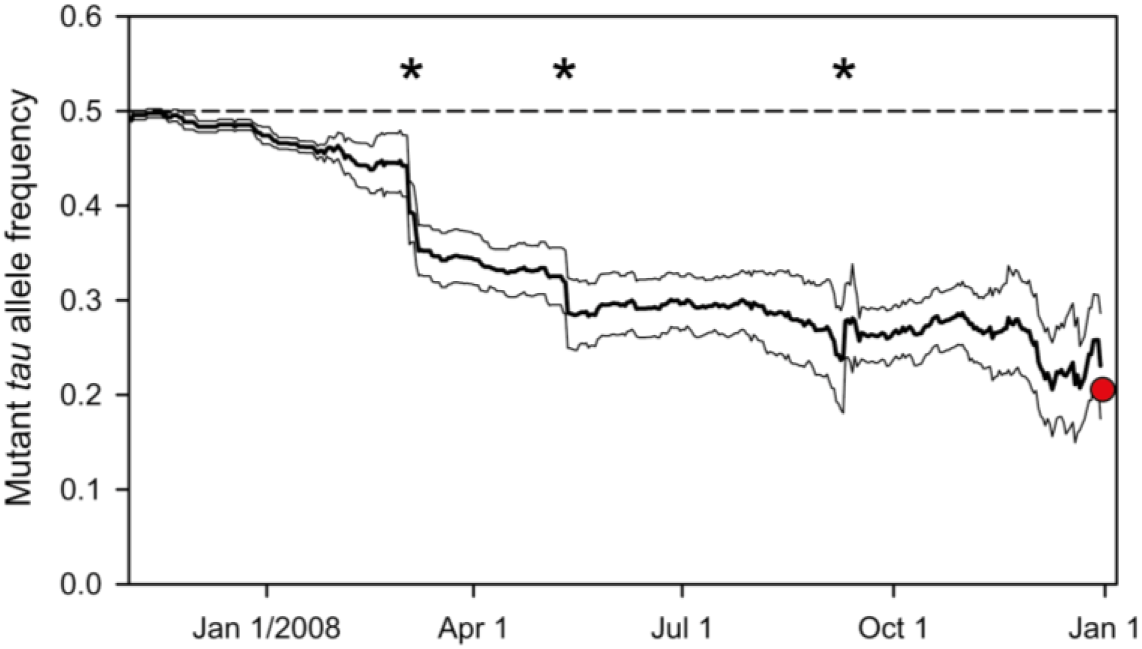
The change of mutant *tau* allele frequency over time. Thick line denotes the mean allele frequency, thin lines its standard errors, dashed line the 50% frequency of mutant allele. Asterisks indicate times at which the offspring of the existing mice were trapped and thus included in the censored population. The red dot indicates the final allele frequency after trapping of all mice at the end of the experiment. At the start, the mutant allele was in near Mendelian ratio (homozygote: heterozygote: wild type = 0.88:2.00:0.78, which resulted in a mutant allele frequency of 49.1%). After a little over a year, the allele frequency had dropped to 20.5%. Figure modified from [15].

These results suggest that fitness consequences of behavioural rhythms with a circadian period length that deviates from the light/dark cycle in a semi-natural setting (second study) may be more severe compared to the consequences of a weaker circadian rhythm (first study). This is in line with the profound impact of strong deviations in circadian period reported from the lab [65,66]. However, more studies on the impact of variation of circadian rhythmicity on fitness in the field are needed.

## Conclusion

So, can we link circadian organization to fitness in the wild? In the second experiment, the ultimate control test would be to shorten the duration of the period of the natural light/dark cycle. However, a true manipulation of the natural light/dark cycle is hard to achieve in the field, and this remains a major limitation for experimental studies on fitness consequences of circadian timing in wild animals. Nevertheless, developing novel, long lasting and smaller tracking systems will expand the possibilities to study natural variation of circadian organization in free-ranging species. These will enable us to follow more and smaller species for a longer time in the field. Indeed, in some contexts (e.g. bees, fish in small ponds, birds), life-long tracking of individuals (e.g. using RFID and satellite tracking; see Case study 1 [38]) is already possible. Information on individual variation in circadian organization, in combination with data on longevity will provide new insights on the evolutionary consequences of daily rhythms in free-ranging animals (Table 1, question 5).

The circadian phenotype of the tracked individuals can be precisely estimated by standard behavioural assays in the laboratory, but also with the use of skin fibroblasts (see Table 3 and [78]). In addition, other manipulations of the natural light/dark cycle (e.g. use of artificial light at night in natural habitat) are possible [79] and have been shown to affect circadian as well as seasonal traits in a variety of species [80–82]. However, the fitness consequences of such effects are still unclear. A recent correlative study has linked light at night to variation in dawn song and reproductive success (extra-pair paternity) in a wild songbird [83], but an experimental manipulation of light at night in the field has shown little effect on the reproductive success in a closely related species [79].

## Outlook

Methods to monitor behavioural, physiological and molecular rhythms develop rapidly, but are not used to their full potential for tracking biologically relevant rhythms in free-ranging animals in their natural environments. Here we have briefly reviewed established and unconventional methods available for tracking animal rhythms in the wild and suggest possible future applications (Table 2 and 3). With the help of three case studies, we further illustrated how to use some of the reviewed technologies to reveal: a) the variability in behavioural rhythms at different taxonomic level, from individuals to species (Case study 1 and 2, Figure 1, 2, 3), (b) the phylogenetic and environmental factors that may influence such variability (Case study 1), and (c) the fitness consequences of functional clocks (Case study 3, Figure 4). These case studies serve as examples of what is currently possible to achieve, but the opportunities do not end here. For instance, whereas detailed tracking data are currently limited to larger animals [45], miniaturization of tags may allow individual tracking of very small animals such as insects [46].

Technological advances not only result in smaller devices, but also ‘smarter’ devices that integrate different sensors. For example, tags that contain accelerometers, as well as physiological and environmental sensors, enable a detailed view into the timing of various animal behaviours and the relation of such timing to habitat characteristics [25,32]. However, one of the challenges is to develop standard methods to extract daily activity patterns from tracking data in order to facilitate intra- and inter-specific comparisons. For example, it is yet unclear how to integrate datasets that differ in resolution (e.g. time interval between fixes, accuracy of location data; but see [11]) or that use different sources of auxiliary data (e.g. accelerometer data compared to instantaneous speed data).

The current rise of field studies is mostly confined to descriptive work. Combining various technologies to record rhythms in the wild with rigorous experimental designs will enhance the mechanistic understanding of how rhythms of free-living animals are regulated, including the relative contribution of endogenous versus environmental factors, as well as the adaptive function of biological clocks.

## Ethics

Research using animals shown in the case studies adhered to local guidelines and appropriate ethical approval and licenses were obtained.

## Data, code and materials

Supporting information, data and R-codes to reproduce Figure 1 and the incubation results of the 1^st^ case study are freely available from Open Science Framework: <u>https://osf.io/wxufm/</u>. Data used to produce Figure 2 will be uploaded in Dryad if the manuscript will be accepted.

## Authors’ contribution

D.M.D., M.B., K.S., S.A. conceived the paper. M.B. drafted the 1st case study, R.K. its foraging part. S.A. drafted the 2^nd^ case study. K.S. drafted the 3^rd^ case study. D.M.D. coordinated the writing and with help from all other authors drafted the introduction, methods review and outlook. All authors finalized the paper and D.M.D. with M.B. addressed referees’ comments.

## Acknowledgments

We thank the co-editor of this special issue Bill Schwarz for guidance during designing and drafting the manuscript, and all that made the data collection and analyses, and this paper possible.

